# Antibody-induced glomerulonephritis is amplified by RTEC-intrinsic IL-17 signaling and restrained by IL-17-mediated induction of the endoribonuclease Regnase-1 (*Zc3h12a*)

**DOI:** 10.1101/2021.01.11.425972

**Authors:** De-Dong Li, Rami Bechara, Kritika Ramani, Chetan V. Jawale, Yang Li, Jay K. Kolls, Sarah L. Gaffen, Partha S. Biswas

**Author notes:** Co-corresponding authors, Partha S. Biswas, S725 BST, Department of Medicine, University of Pittsburgh, 200 Lothrop Street, Pittsburgh, PA 15261, Sarah L. Gaffen, S702 BST, Department of Medicine, University of Pittsburgh, 200 Lothrop Street, Pittsburgh, PA 15261.

## Abstract

Antibody-mediated glomerulonephritis (AGN) is a clinical manifestation of many autoimmune kidney diseases for which few effective treatments exist. Chronic inflammatory circuits in renal glomerular and tubular cells lead to tissue damage in AGN. These cells are targeted by the cytokine IL-17, which has recently been shown to be a central driver of the pathogenesis of AGN. However, surprisingly little is known about the regulation of pathogenic IL-17 signaling in the kidney. Here, using a well characterized mouse model of AGN, we show that IL-17 signaling in renal tubular epithelial cells (RTECs) is necessary for AGN development. We also show that Regnase-1, an RNA binding protein with endoribonuclease activity, is a negative regulator of IL-17 signaling in RTECs. Accordingly, mice with a selective Regnase-1 deficiency in RTECs exhibited exacerbated kidney dysfunction in AGN. Mechanistically, Regnase-1 inhibits IL-17-driven expression of the transcription factor IκBξ and consequently its downstream gene targets including *Il6* and *Lcn2*. Moreover, deletion of Regnase-1 in human RTECs reduced inflammatory gene expression in an IκBξ-dependent manner. Overall, these data identify an IL-17-driven inflammatory circuit in RTECs during AGN that is constrained by Regnase-1.

## Introduction

Antibody-mediated glomerulonephritis (AGN) describes a heterogenous group of renal conditions caused by an inappropriate response to renal autoantigens such as the glomerular basement membrane (GBM) (1, 2). The most severe form of human AGN is crescentic GN, characterized by the formation of glomerular crescents and tubulointerstitial inflammation (3, 4). Although the initiators of AGN differ among diseases, the terminal events in end-organ kidney damage share common mechanisms that remain poorly understood. Even with the most aggressive immunosuppressive regimens, as many as ~30% of nephritic patients develop end-stage renal disease (5). To date, the best therapies are broad-acting immunosuppressants, namely cyclophosphamide with corticosteroids (5). Gaining a fundamental understanding of how pathogenic kidney inflammation is driven and amplified could help fill an unmet clinical need to achieve more specific and effective responses in AGN.

Typically, immune-mediated tissue pathology is viewed through the lens of hematopoietic lymphoid and myeloid cells. Less appreciated is the contribution of organ-specific tissue cells as amplifiers of inflammation that ultimately lead to chronic disease. In the kidney, glomerular and tubular inflammation are hallmark features of AGN (6). In fact, the majority of AGN patients progress to end-stage renal disease as a consequence of chronic tubular inflammation (7). Renal tubular epithelial cells (RTECs) comprise the major cell type of the tubular compartment (8). These cells actively contribute to the production of inflammatory mediators that propagate tissue injury locally by secreting cytokines and chemokines and undergoing apoptosis (7). Even so, our understanding of the inflammatory circuits operative in RTECs during chronic kidney disease is surprisingly rudimentary.

AGN has traditionally been considered a B-cell-dependent disease because of the high autoantibody levels found in these patients (9). However, multiple studies also implicate IL-17, the signature cytokine of Th17 cells, as an essential driver of AGN pathology (10–15). IL-17 fuels an inflammatory response that damages renal podocytes, causing proteinuria. Loss of glomerular filtration is followed by an IL-17-dominated tubular inflammation and damage of RTECs. Interestingly, IL-17 signaling in the nephritic kidney has no impact on the immune-complex deposition in the glomeruli (10). While IL-17 is secreted almost exclusively by lymphocytes, its target cells are often but not always non-hematopoietic (16, 17). The specific IL-17 target cells in the context of renal autoimmunity remain unclear.

Although not well defined in the kidney, dysregulation of Th17 cells or IL-17 signaling causes pathology in other autoimmune conditions, such as psoriasis and psoriatic arthritis, where anti-IL-17 biologics are now standard therapies (18, 19). A landmark recent study showed that human Th17 cells specific for commensal microbes such as *Staphylococcus aureus* and *Candida albicans* contribute to AGN (20). These findings are consistent with prior work in preclinical models that IL-17 and its adaptor Act1 promote AGN in mice (10–14, 21–23). The downstream events operative in AGN have been presumed to occur through IL-17-mediated activation of the NF-κB signaling pathway, which is induced in response to a myriad of immune stimuli, including IL-17, TNF, TLR ligands and T cell signaling (24, 25). NF-κB induces a panoply of genes that promote renal inflammation, including IL-6, neutrophil-attracting chemokines (CXCL1, CXCL5) and lipocalin 2 (Lcn2, NGAL, 24p3), a major biomarker and driver of renal inflammation (26–28). NF-κB also upregulates the noncanonical NF-κB family member IκBξ (*Nfkbiz*), a transcription factor that regulates a more restricted subset of genes that includes both IL-6 and Lcn2 (29, 30). In recent years it has become evident that IL-17 regulates many genes post-transcriptionally, either by controlling mRNA stabilization and/or by influencing translation of downstream transcripts (31–33). Among these post-transcriptionally regulated targets are *Il6* and *Nfkbiz* (IκBξ).

Autoimmune signaling must be sufficiently restrained in order to prevent hyper-activation of pathways that could contribute to immune-mediated pathology. The endoribonuclease Regnase-1 (a.k.a. MCPIP1, encoded by *Zc3h12a*) is best recognized as an inhibitor of Toll-like receptor (TLR) and TCR signaling (34). In CD4^+^ T cells, loss of Regnase-1 leads to markedly elevated IL-6 levels, and consequently, elevated Th17 differentiation and exacerbated autoimmune disease, demonstrated in the experimental autoimmune encephalitis (EAE) model of multiple sclerosis (35). In cultured fibroblasts, Regnase-1 has been shown to restrict IL-17 signaling by degrading downstream target mRNAs such as *Il6* (33). With respect to kidney, however, little is known, though a global deficiency of Regnase-1 enhances the immune response to disseminated *C. albicans*, a fungal infection that targets the kidney (33).

In this report, we show that RTECs are critical mediators of inflammatory damage in AGN by virtue of their sensitivity to IL-17 signaling. Conversely, Regnase-1 restricts AGN pathogenesis by limiting IL-17-driven renal-specific expression of the transcription factor IκBξ and consequently its downstream gene targets *Il6* and *Lcn2* within RTECs. These findings demonstrate a potent, kidney-intrinsic inflammation circuit that is potentiated by RTEC-specific IL-17 signaling and restrained by Regnase-1 in a highly cell-type specific manner.

## Results

### RTEC-intrinsic IL-17 signaling is critical for AGN pathology

AGN is mediated by both kidney-infiltrating immune cells (hematopoietic) and kidney-resident non-hematopoietic cells. Several studies show that IL-17 and its receptor (IL-17RA), through the adaptor Act1, are essential for AGN (10–14, 21). However, the identity of IL-17 target cells in this setting is unknown. To define the contribution of IL-17RA in hematopoietic *vs*. non-hematopoietic cells, we created bone marrow (BM) chimeric mice in which *Il17ra^-/-^* or wild type (WT) BM was adoptively transferred into irradiated reciprocal hosts. Successfully reconstituted recipients were then evaluated for susceptibility to AGN by injecting mice with rabbit IgG in Complete Freund’s adjuvant (CFA), followed three days by injection with rabbit anti-GBM serum (10, 36). As shown, WT hosts receiving *Il17ra^-/-^* BM showed identical susceptibility to AGN as WT mice, indicating that any activity of IL-17 in hematopoietic cells is dispensable for AGN pathology. Conversely, *Il17ra^-/-^* mice, regardless of BM source, were resistant to AGN (Fig 1A). Thus, IL-17RA signaling in non-hematopoietic cells is required for development of AGN.

**Fig 1:**
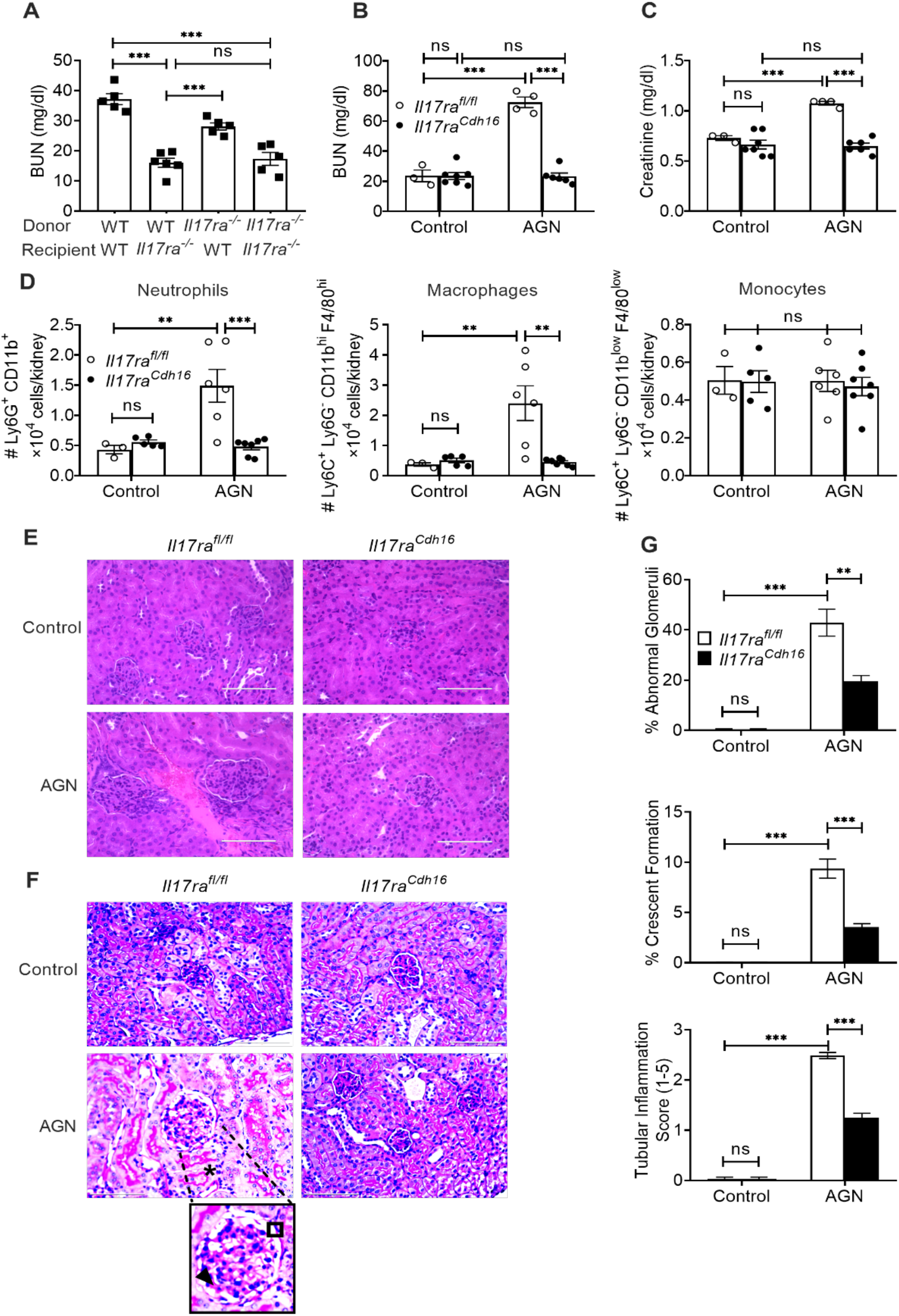
IL-17RA signaling in RTECs is required for AGN. **(A)** Bone marrow (BM) cells from *Il17ra^−/−^* (CD45.2^+^) and wild type (WT) (CD45.1^+^) mice were adoptively transferred into sub-lethally irradiated *Il17ra^−/−^* or WT recipients (n=5-6). Eight weeks later, successfully reconstituted mice were subjected to AGN and assessed for kidney dysfunction by measuring serum BUN levels. *Il17ra^fl/fl^* and *Il17ra^Cdh16^* mice (*n*=3-7) were subjected to AGN. At day 14 post anti-GBM serum injection, **(B)** serum BUN, and **(C)** serum creatinine levels were measured by ELISA. **(D)** Neutrophils, macrophages and monocytes infiltration in the kidney was quantified by flow cytometry at day 7 p.i. Representative photographs of H&E-stained **(E)** and PAS-stained **(F)** renal histopathology were assessed. **(G)** Renal pathology was blindly evaluated and scored for percentages of abnormal glomeruli and crescent formation and tubular inflammation score. Data representative of 1 of 3 mice/group for E and F. Magnification: 400X. Open square: indicating entire glomerulus with mesangial and endocapillary hypercellularity; black arrow: GBM thickening; *: tubular atrophy. Data pooled from at least 2 independent experiments. Statistical analysis by One-way ANOVA (A) and Two-way ANOVA (others).

The kidney-intrinsic non-hematopoietic cells thought to mediate AGN are RTECs. Prior studies showed that IL-17 can act on primary cultured RTECs to induce expression of cytokine and chemokine genes that are known to drive inflammation in AGN (24). Based on this, we sought to define the role of IL-17 signaling in RTECs *in vivo*. We took advantage of a well characterized *Cdh16^Cre^* mice, in which the Cre recombinase is expressed in the epithelial lining of proximal tubules, collecting ducts, loops of Henle and distal tubules (37). We crossed *Il17ra^fl/fl^* mice to *Cdh16^Cre^* mice (termed *Il17ra^Cdh16^*), which selectively and efficiently depleted IL-17RA in RTECs, as previously reported (38). Following AGN, *Il17ra^Cdh16^* mice showed diminished levels of serum BUN and creatinine (Fig 1B and C). *Il17ra^Cdh16^* kidneys exhibited compromised renal infiltration of neutrophils and macrophages at day 7 post-AGN (Fig 1D and Fig S1). Interestingly, the number of inflammatory monocytes, CD4^+^ T, CD8^+^ T and B cells were comparable between the groups (Fig 1D and Fig S1 and S2A-B). When evaluated for kidney pathology, *Il17ra^Cdh16^* kidneys showed reduced mesangial and endocapillary hypercellularity, thickening of GBM, crescents formation, tubular atrophy and tubular inflammatory cells infiltration in comparison to controls, similar to *Il17ra^-/-^* or *Act1^-/-^* mice (10, 14) (Fig 1E-G). These results thus demonstrate an RTEC-specific role of IL-17R signaling in AGN.

### Regnase-1 haploinsufficient mice show exaggerated AGN

Surprisingly little is known about the molecular signals that constrain inflammation in the context of AGN. Regnase-1 (encoded by *Zc3h12a*) is a negative feedback regulator of many inflammatory stimuli, including the TCR, TLR4 and IL-17 (33, 34). Mice lacking *Zc3h12a* entirely show systemic inflammation and early mortality, an effect that we previously found to be independent of IL-17 (39, 40). In contrast, *Zc3h12a* haploinsuffìcient (*Zc3h12a*^+/-^) mice have no peripheral inflammation in kidney or other visceral organs but exhibit enhanced susceptibility to IL-17-induced pathology in certain settings, including EAE (33, 41, 42). Consequently, we subjected *Zc3h12a^+/-^* mice to AGN to determine if this molecule influences outcomes in renal autoimmunity. As shown, *Zc3h12a*^+/-^ mice were markedly more susceptible to AGN than control mice (Fig 2A). Mice lacking single copy of *Zc3h12a* gene demonstrated increased infiltration of neutrophils and macrophages but not monocytes, CD4^+^ T, CD8^+^ T and B cells (Fig 2B and Fig S1 and S3A-C). Moreover, kidney dysfunction was reversed in *Zc3h12a^+/-^Il17ra^-/-^* mice, indicating that IL-17 signaling is essential for Regnase-1-mediated inhibitory signaling in the nephritic kidney (Fig 2C).

**Fig 2:**
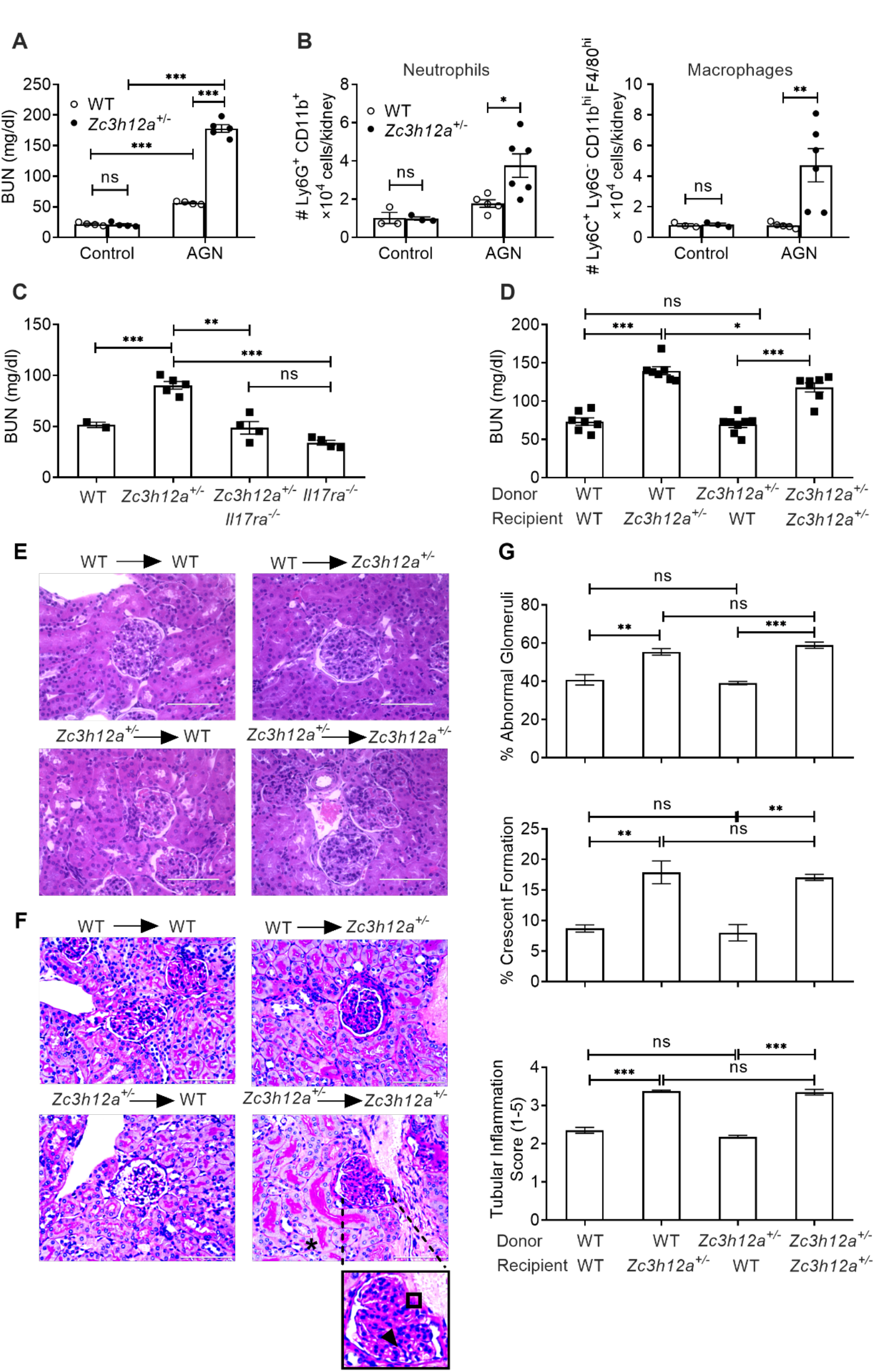
Zc3h12a^+/-^ mice show exaggerated AGN. **(A)** *Zc3h12a^+/+^* (WT) and *Zc3h12a^+/-^* (n=4-5) were subjected to AGN. Mice were evaluated for kidney dysfunction by measuring serum BUN at day 14 p.i. **(B)** Neutrophils and macrophages infiltration in the kidney was quantified by flow cytometry at day 7 p.i. **(C)** WT, *Zc3h12a^+/-^, Zc3h12a^+/-^Il17ra^-/-^ and Il17ra^-/-^* (n=3-5) were subjected to AGN. Serum BUN level was measured at day 14 p.i. **(D)** BM cells from *Zc3h12a^+/−^* and WT mice were adoptively transferred into sub-lethally irradiated *Zc3h12a^+/−^* or WT recipients (n=5-7). Eight weeks later, successfully reconstituted mice were subjected to AGN and assessed for serum BUN levels. Representative photographs of **(E)** H&E-stained and **(F)** PAS-stained renal histopathology. Data representative 1 of 3 mice/group. Magnification: 400X. Open square: indicating entire glomerulus with mesangial and endocapillary hypercellularity; black arrow: GBM thickening; *: tubular atrophy. **(G)** Renal pathology was blindly evaluated and scored for percentages of abnormal glomeruli and crescent formation and tubular inflammation score. Data pooled from at least 2 independent experiments. Statistical analysis by Two-way ANOVA (A and B) and One-way ANOVA (C, D and G).

Regnase-1 can also restrain Th17 cell differentiation, and hence mice lacking Regnase-1 in CD4^+^ T cells have exacerbated disease in EAE (35). Thus, the increased nephritis in *Zc3h12a^+/-^* could be due to enhanced IL-17 production from T cells or to unrestrained IL-17 signaling in responder cells, or potentially both. To distinguish among these possibilities, we created BM chimeric mice where *Zc3h12a^+/-^* or littermate *Zc3h12a^+/+^* BM were adoptively transferred into irradiated reciprocal hosts. Successfully reconstituted mice were subjected to AGN. These data show that Regnase-1 in non-hematopoietic cells was required for AGN development, evidenced by increased serum BUN and creatinine (Fig 2D and S3D). *Zc3h12a^+/-^* animals receiving either WT or *Zc3h12a^+/-^* BM showed evidence of increased glomerular pathology as characterized by exaggerated crescents formation, mesangial hypercellularity and increased GBM thickness. Additionally, these animals exhibited increased tubular atrophy and tubular inflammatory cells infiltration than WT recipients (Fig 2E-G). These data indicated that Regnase-1 restricts AGN pathogenesis by acting in non-hematopoietic cells, though with the caveat that mice still retained one intact copy of the *Zc3h12a* gene.

### Regnase-1 restrains AGN severity in an RTEC-intrinsic manner

Given that Regnase-1 restricts IL-17 signaling and that Regnase-1 and IL-17RA both act in non-hematopoietic cells in AGN in an opposing manner, we predicted that loss of Regnase-1 in RTECs would exacerbate tissue pathology in AGN. We crossed *Zc3h12a^fl/fl^* and *Cdh16^Cre^* mice to delete Regnase-1 in RTECs (*Zc3h12a^Cdh16^*). Mice lacking Regnase-1 in RTECs breed normally and displayed normal serum BUN and kidney tissue architecture at baseline (Fig 3A and D). After AGN induction, compared to *Zc3h12a^fl/fl^* controls, *Zc3h12a^Cdh16^* mice showed more severe kidney dysfunction and a trend toward increased expression of the classical kidney injury marker *Kim1* (43) (Fig 3A-C). Moreover, *Zc3h12a^Cdh16^* mice exhibited more glomerular abnormality as characterized by increased mesangial and endocapillary hypercellularity, thickening of GBM and crescents formation following AGN (Fig 3D-F). *Zc3h12a^Cdh16^* kidneys demonstrated exaggerated tubular atrophy and inflammatory cell infiltration in the tubular space than control mice. *Zc3h12a^Cdh16^* kidney also showed increased expression of multiple chemokines implicated in AGN including *Cxcl1, Cxcl2, Ccl20* and *Ccl2* that are known to recruit immune cells to diseased tissue (Fig 4A). Consequently, *Zc3h12a^Cdh16^* mice exhibited a trend of increased kidney-infiltrating CD45^+^ inflammatory cells (Fig S1 and S4A). Although the number of neutrophils and macrophages were increased in *Zc3h12a^Cdh16^* mice, the frequencies of monocytes, CD4^+^ T, CD8^+^ T and B cells were comparable among the nephritic groups (Fig 4B and C, Fig S1 and S4B). These results indicate a role for Regnase-1 as a negative regulator of renal inflammation.

**Fig 3:**
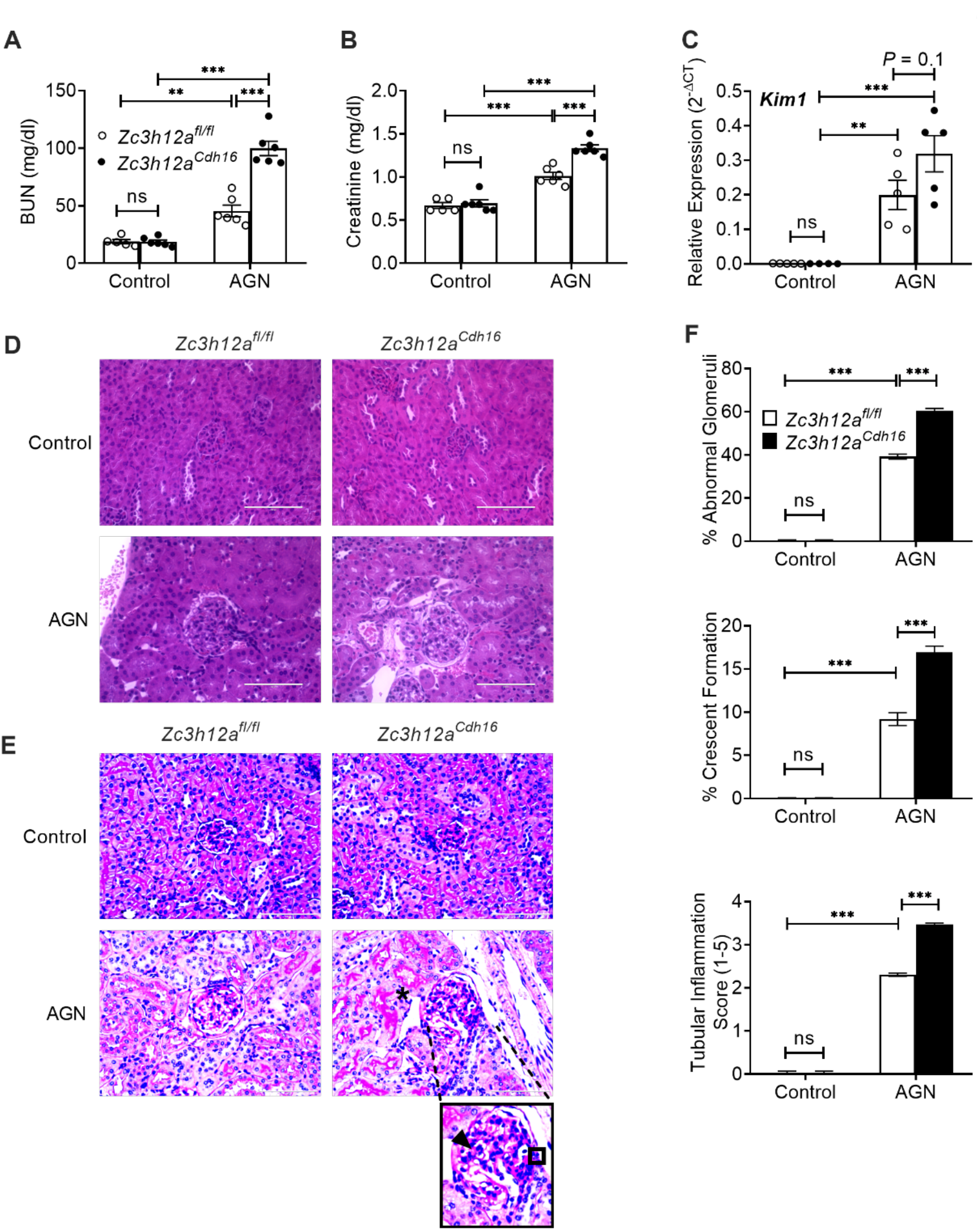
RTEC-specific Regnase-1 deficiency exacerbates AGN. *Zc3h12a^fl/fl^* and *Zc3h12a^Cdh16^* mice (*n*=5-6) were subjected to AGN. At day 14 p.i., **(A)** serum BUN, and **(B)** serum creatinine level were assessed. **(C)** *Kim1* transcript expression was assessed at day 7 p.i. Expression was normalized to *Gapdh*. Representative photographs of **(D)** H&E-stained and **(E)** PAS-stained renal histopathology. **(F)** Renal pathology was blindly evaluated and scored for percentages of abnormal glomeruli and crescent formation and tubular inflammation score. Data representative of 1 of 3 mice/group for D and E. Magnification: 400X. Open square: indicating entire glomerulus with mesangial and endocapillary hypercellularity; black arrow: GBM thickening; *: tubular atrophy. Data pooled from at least 2 independent experiments. Statistical analysis by Two-way ANOVA.

**Fig 4:**
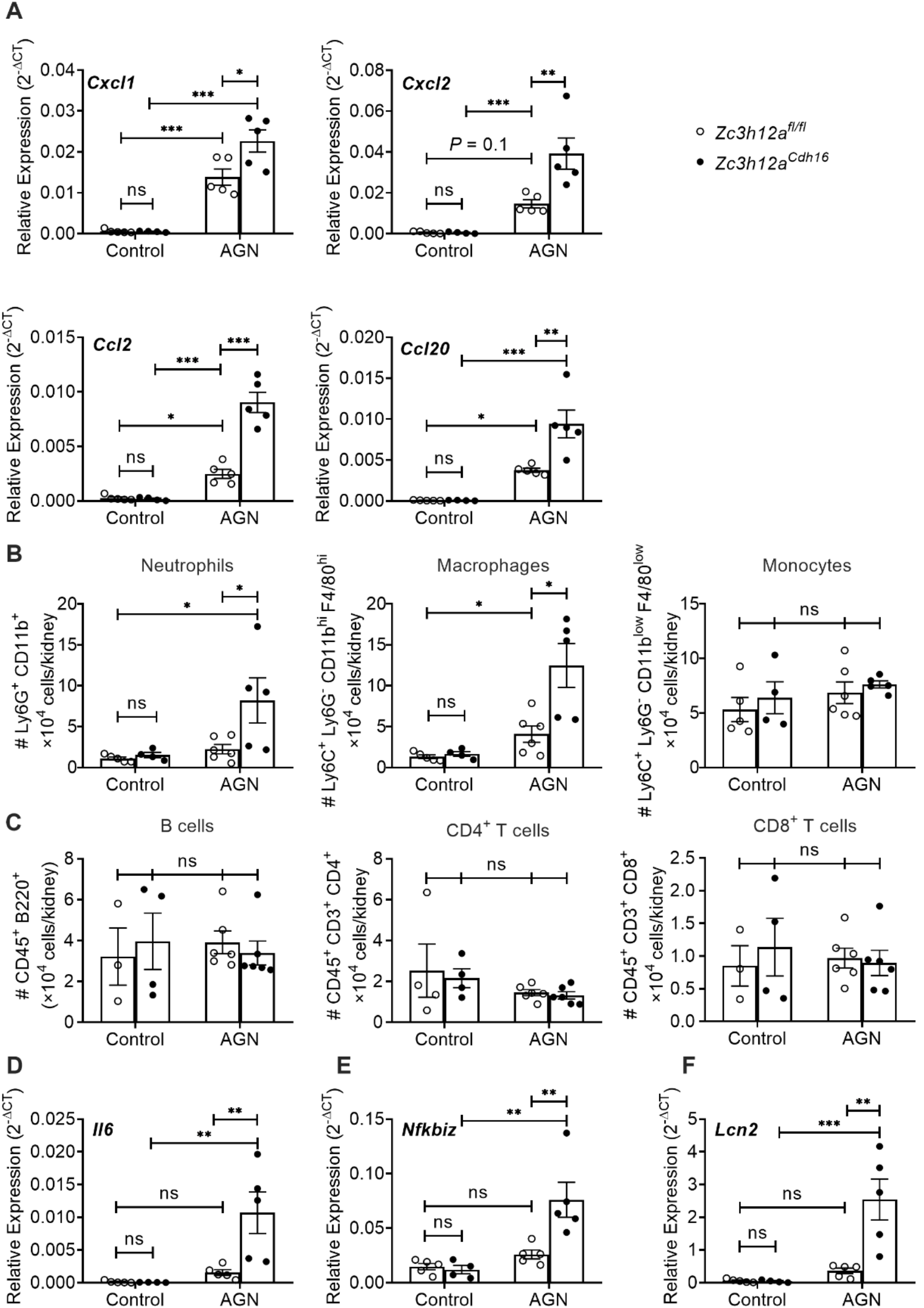
Increased inflammatory gene expression and cell infiltration in the nephritic kidney of RTEC-specific Regnase-1 deficient mice. *Zc3h12a^fl/fl^* and *Zc3h12a^Cdh16^* mice (*n*=3-6) were subjected to AGN. At day 7 p.i., **(A)** renal *Cxcl1, Cxcl2, Ccl20* and *Ccl2* mRNA expression was measured by qPCR, **(B, C)** neutrophils, macrophages, monocytes, B cells, CD4^+^ T cells and CD8^+^ T cells infiltration in the kidney was quantified by flow cytometry, and **(D)** *Il6*, **(E)** *Nfkbiz*, and **(F)** *Lcn2* transcript expression was evaluated by qPCR. Expression was normalized to *Gapdh* (A, D, E and F). The data is pooled from at least 2 independent experiments. Statistical analysis by Twoway ANOVA.

Regnase-1 functions by binding to and degrading client mRNA transcripts (33). Some inflammatory transcripts are direct targets of Regnase-1 such as *Il6*, where Regnase-1 binds to a stem-loop sequence in *Il6* the 3’ UTR to promote degradation. However, many mRNAs are indirectly influenced by Regnase-1 through its capacity to degrade mRNA including the gene encoding the IκBξ transcription factor (*Nfkbiz*). A major example of IκBξ-dependent mRNA is *Lcn2* (44). We therefore examined expression of known Regnase-1 client transcripts known to be involved in AGN pathology in *Zc3h12a^Cdh16^* mice. There was no difference in the expression of *Il6*, *Nfkbiz* or *Lcn2* in kidney at baseline (Fig 4D-F). However, following AGN, there was a marked increase in levels of *Il6*, *Nfkbiz* and *Lcn2* in *Zc3h12a^Cdh16^* kidneys compared to controls. Notably, all these genes are also known targets of IL-17 in AGN and are reduced in *Il17ra^-/-^* mice subjected to AGN (10). Collectively, these data suggest a role for Regnase-1 in restraining expression of pathogenic IL-17-dependent genes in an RTEC-specific manner during the inflammatory process of AGN.

### Regnase-1 in human RTECs drives inflammation via IκBξ

To determine in a more direct way how Regnase-1 functions in RTECs, we deleted the *ZC3H12A* gene in the human tubular kidney 2 (HK-2) cell line using CRISPR-Cas9 technology (henceforth referred to as *HK-2^ΔZC3H12A^*) (Fig 5A and B). HK-2 is a tubular epithelial cell line derived from normal kidney, that we find to be highly responsive to IL-17 (Fig 5C). HK-2^*ΔZC3H12A*^ and control HK-2 cells were stimulated with IL-17 and assessed for mRNA and protein expression of known IL-17 responsive genes including *IL6*, *LCN2, NFKBIZ, CEBPB* and *CEBPD*. We also performed stimulations in conjunction with TNFa, since IL-17 and TNFa often signal in a potently synergistic manner (45). IL-17 alone or in conjunction with TNFα upregulated transcript (Fig 5C and E) and protein (Fig 5D and F) expression of *IL6* and *LCN2* in HK-2^*ΔZC3H12A*^ cells. Notably, HK-2^*ΔZC3H12A*^ cells demonstrated an increased expression of *NFKBIZ* transcript and IκBξ expression following IL-17 treatment (Fig 5G and H). The impact of IL-17 alone or IL-17 + TNFα on the mRNA and protein expression of *CEBPB* and *CEBPD* was variable (Fig S5A and B). Additionally, luciferase assay revealed activation of *Lcn2* promoter activity following IL-17 stimulation in human embryonic kidney 293 cells (HEK-293) (Fig S5C). Overall, these results indicate that Regnase-1 is a negative regulator of IL-17 signaling in human RTECs.

**Fig 5:**
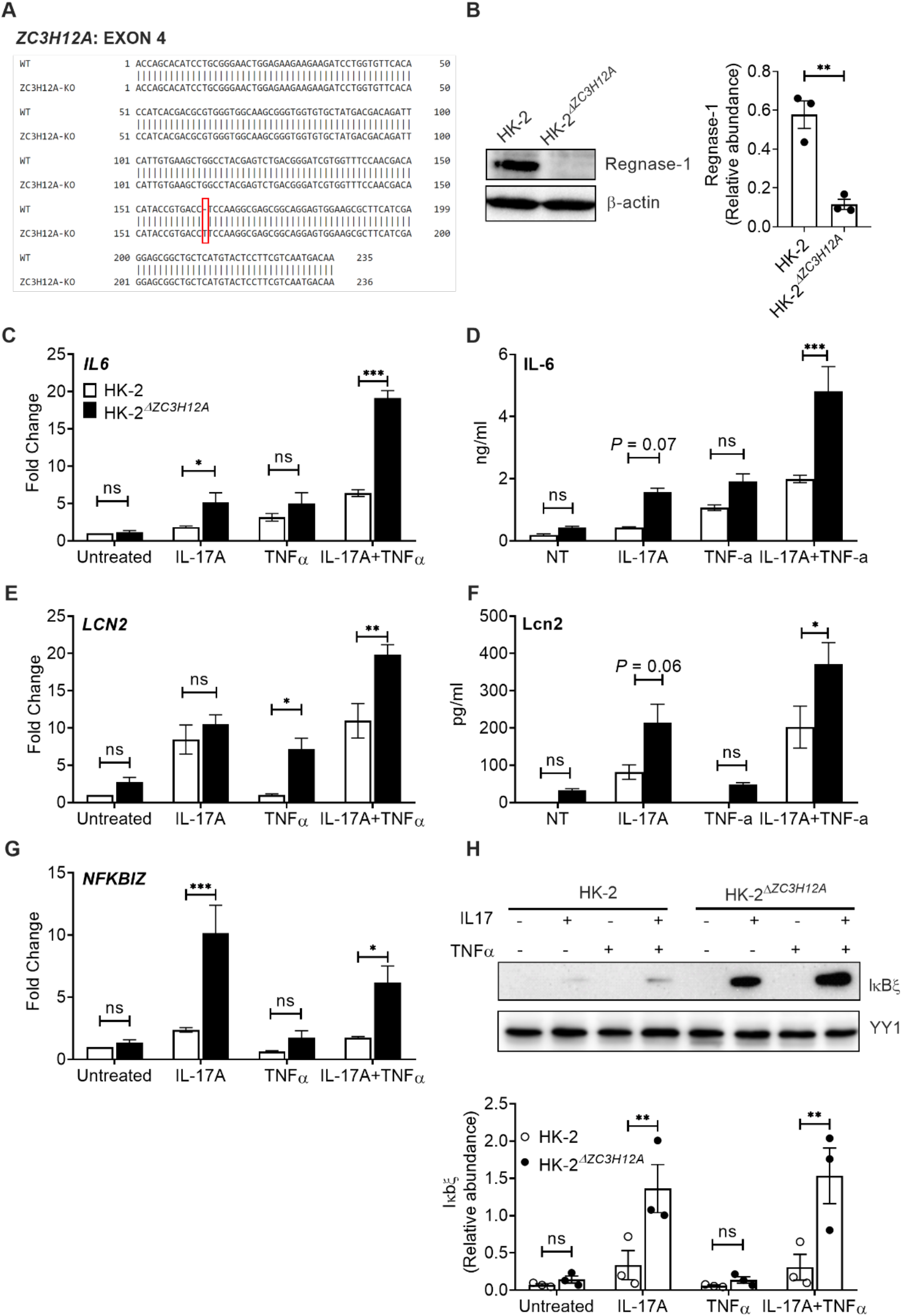
Deletion of ZC3H12A gene elevates inflammatory mediator expression in the HK-2 cell line. **(A)** Experimental strategy to delete ZC3H12A gene by CRISPR-Cas9 in the HK-2 cell line. (B) HK-2 and HK-2^*ΔZC3H12A*^ cells were lysed and evaluated for Regnase-1 by western blot. Protein relative abundance and representative image are shown. HK-2 and HK-2^*ΔZC3H12A*^ cells were stimulated with IL-17 and/or TNFa and (C, E, G) mRNA expression of IL6, LCN2 and NFKBIZ was measured by qPCR (8 h post-stimulation), normalized to GAPDH. (D, F) Protein level of IL-6 (8 h post-stimulation) and Lcn2 (24 h post-stimulation) in culture supernatant were measured by ELISA. (H) HK-2 and HK-2^*ΔZC3H12A*^ cells were stimulated with IL-17 and/or TNFa for 8 h and cell lysates were evaluated for IκBξ by western blot. Protein relative abundance and representative image are shown. Data pooled from at least 3 independent experiments. Statistical analysis by unpaired T test (B) or Two-way ANOVA (others).

Since Regnase-1 restricts expression of the IκBξ transcription factor (*NFKBIZ*), which mediates expression of many of these characteristic downstream mRNAs, we evaluated the impact of IκBξ in HK-2 cells with respect to IL-17 signaling. To that end, we knocked down *NFKBIZ* by siRNA in HK-2^*ΔZC3H12A*^ and control HK-2 cells, which resulted in 50-60% reduction in *NFKBIZ* expression (Fig 6A). HK-2^*ΔZC3H12A*^ cells in which *NFKBIZ* was silenced showed diminished levels of *LCN2* and *IL6* mRNA compared to controls (Fig 6B). Interestingly, while *CCL20* levels were reduced upon *NFKBIZ* silencing, *CXCL1* and *CXCL2* were increased (Fig 6C). These results indicate that Regnase-1 selectively suppresses some IL-17 responsive genes that mediate kidney inflammation by increasing *NFKBIZ* in human RTECs (Fig 7).

**Fig 6:**
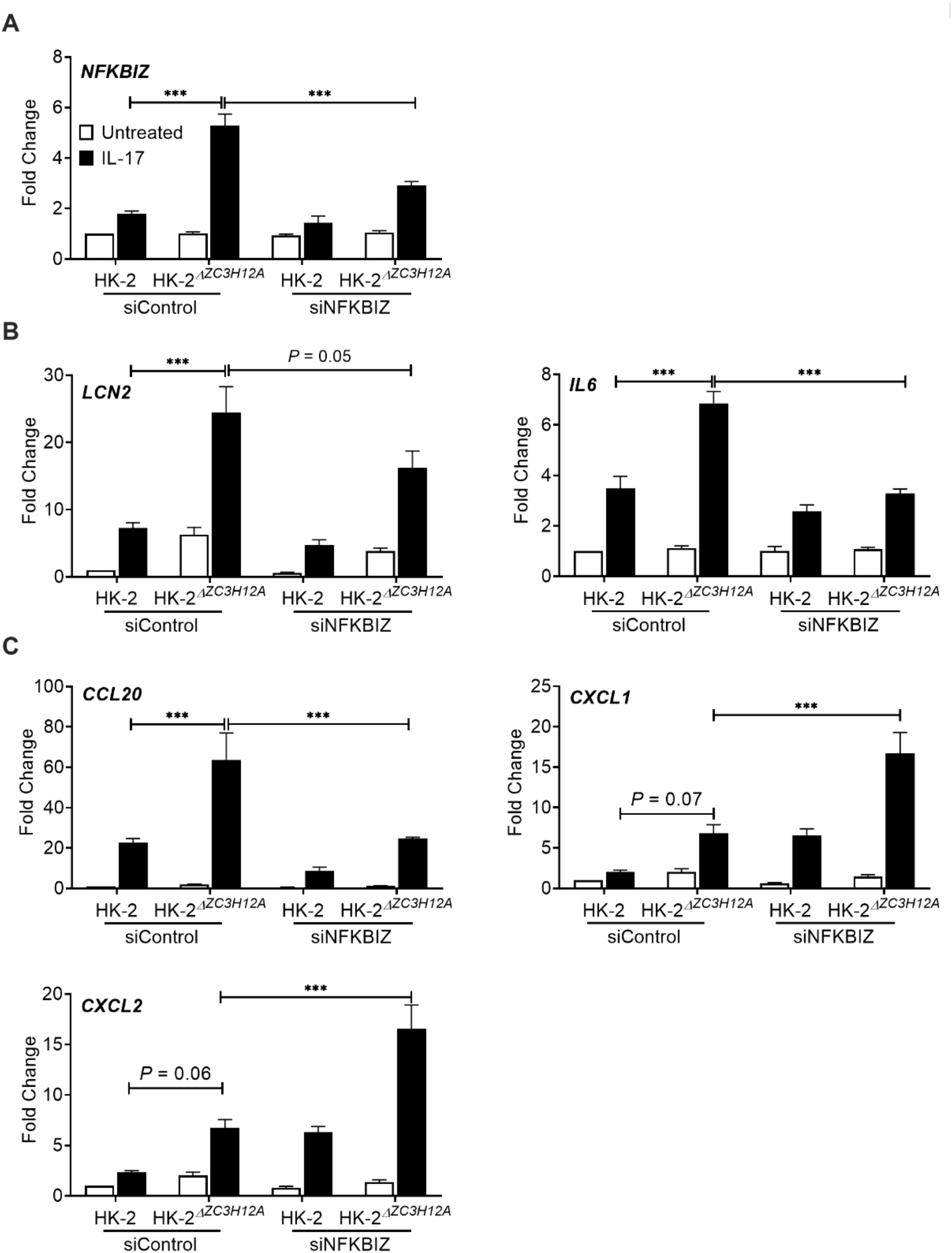
Regnase-1 upregulates IL6 and LCN2 gene expression via NFKBIZ. HK-2 and HK-2^*ΔZC3H12A*^ cells were either treated with siRNA against *NFKBIZ* or control siRNA. **(A)** Efficiency of *NFKBIZ* gene silencing was measured by qPCR. **(B)** Cells were either stimulated with IL-17 or left untreated for 8 h and analyzed for *IL6, LCN2*, and **(C)** *CCL20, CXCL1* and *CXCL2* gene expression by qPCR. Data pooled from at least 3 independent experiments. Statistical analysis by Two-way ANOVA.

**Fig 7:**
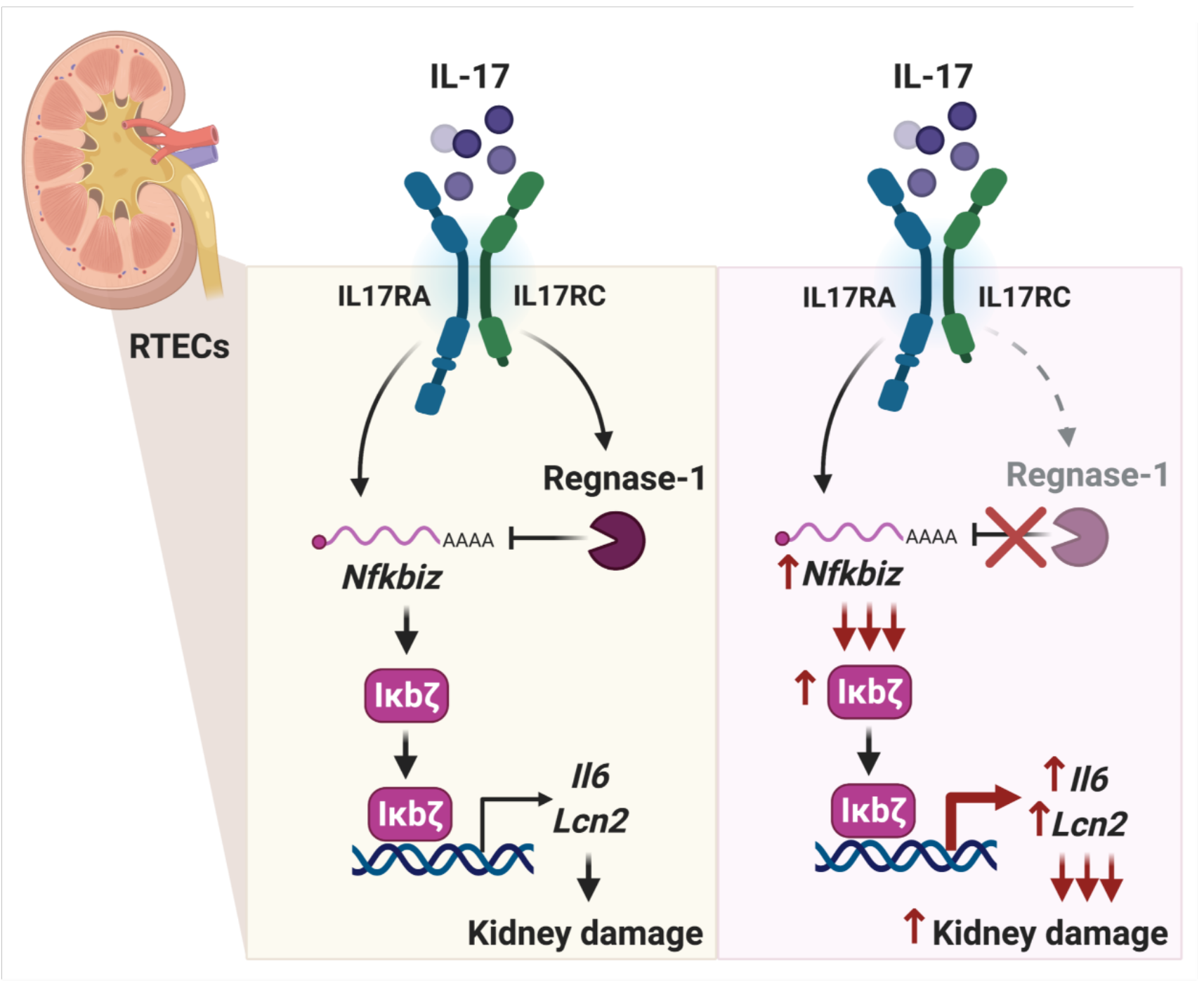
RTEC-specific pathogenic IL-17 signaling is restrained by Regnase-1. IL-17 binds to its receptor (IL-17RA) on RTECs and transcriptionally activates transcription factors such as IκBζ (*Nfkbiz*). These transcription factors turn on genes such as *Il6* and *Lcn2*, which promote kidney inflammation and damage in AGN. IL-17 signaling also induces a pathway that destabilizes mRNA through the endoribonuclease Regnase-1, which binds to the 3’ UTR of target mRNAs. One key target of Regnase-1 is *Nfkbiz*. Hence, Regnase-1 restrains pathogenic IL-17 signaling in RTECs and consequently AGN pathology by negatively regulating the expression of IκBζ responsive genes.

## Discussion

Development of improved therapeutic approaches to treat AGN will require a more comprehensive understanding of the fundamental immune mechanisms in the kidney that drive this disease. The kidney is an immunologically distinct organ and is exquisitely sensitive to endorgan damage in numerous autoimmune diseases and infections. Indeed, renal injury caused by inappropriate immune responses is the most important predictor of mortality in AGN patients (7). Even so, our understanding of the fundamental immune processes in the kidney lags considerably behind that of other visceral organs such as the gut or liver (1, 46). This is due to multiple kidney-specific factors including poor regenerative capacity of nephrons, uremic toxins, hypoxia and arterial blood pressure, which have confounding impacts on the ongoing immune response in the kidney. Defining how IL-17 mediates signals in target cell types in the nephritic kidney has important therapeutic implications, as specific targeting of the IL-17 pathway in these kidney-resident cells could spare other organs from unwanted side effects and potentially preserve important IL-17 host-defensive activities.

IL-17 is a critical contributor of autoinflammatory diseases (47). Antibodies that neutralize IL-17 were approved in 2016 and show efficacy in psoriasis, psoriatic arthritis and ankylosing spondylitis (18, 19). In contrast, IL-17 blockade caused adverse outcomes in Crohn’s disease due to an unexpectedly strong tissue-reparative effect of intestinal epithelium (48–50). These dichotomous outcomes highlight the ramifications of tissue-specific mechanisms of IL-17 signaling. While there has been much research effort describing IL-17-producing cells and their generation, far less is known about how IL-17 mediates downstream signal transduction and mechanisms that regulate this process in responding cells. We showed that RTEC-specific activity of the IL-17 receptor subunit IL-17RA drives chronic inflammatory response in the nephritic kidney. Notably, IL-17RA acts as a shared receptor subunit for the related cytokines IL-17F and IL-17C (51). Hence, some of the effects seen here could be attributable to these cytokines as well. Despite their emerging role in AGN, both IL-17F and IL-17C signaling mechanisms in the RTECs are almost entirely undefined, and no studies to date have focused on the regulation of IL-17F and IL-17C-dependent signal transduction in these kidney-resident cell types (52, 53). Moreover, the type 2 cytokine IL-25 also signals via IL-17RA (51). To date, there is no evidence suggesting a pathogenic role of IL-25 in AGN development. The findings in this study thus set the stage for pursuing analysis of these related but poorly understood IL-17-family cytokines.

The contribution of kidney-resident cells such as RTECs is often overlooked when considering the development of new therapeutic strategies in inflammatory kidney diseases. Although the IL-17R is ubiquitously expressed (51) and some studies indicated that IL-17 can act on immune cells including T cells, B cells and NK cells (16, 17, 54), we show that the contribution of the hematopoietic system in AGN is in fact negligible. Rather, IL-17, produced locally in the nephritic kidney by Th17 and γδ T cells, signals directly on RTECs to drive production of pathogenic factors including cytokines, chemokines and Lcn2 (10). This finding is in line with other organ-specific autoimmune conditions including collagen-induced arthritis and imiquimod (IMQ)-induced dermatitis where a tissue-dependent role for IL-17 signaling was similarly demonstrated. A limitation of BM chimeric mouse studies is that kidney harbors a radio-resistant tissue-resident population of macrophage (CX3CR1^+^), which are responsive to IL-17 (55). Hence, future studies will need to address the role of pathogenic IL-17 signaling in tissue-resident macrophages in AGN, e.g., by crossing *CX3CR1^Cre^* mice to *Il17ra^fl/fl^* animals.

Regnase-1 has been shown to restrict many inflammatory signals, and IL-17 is just one of many. Based on our studies, we cannot rule out the roles for Regnase-1 in restricting other tissue-dependent inflammatory signals that use the IL-17RA subunit, including IL-17F, IL-17C, IL-25 and non-IL-17 cytokines such as IL-1 family members. Since Regnase-1 acts on T cells and macrophages, it is also surprising that our BM chimera and RTEC-specific deletion studies indicated that the impact of Regnase-1 in AGN seemed to be so RTEC-specific (35, 56). However, this finding does suggest that a major function of Regnase-1 is to restrain downstream IL-17R-mediated signals in the kidney. We previously reported that mice with a Regnase-1 deficiency have increased resistance to kidney infections caused by systemic *Candida albicans* infections (33). These data indicate that, while constraint of IL-17 signaling by Regnase-1 is beneficial in preventing autoimmunity, this same inhibition limits the efficacy of IL-17 in mediating renal host defense, and thus care will be needed when considering targeting this pathway pharmacologically. Although Cre-lox technology has enabled manipulation of gene expression in targeted tissues, off-target effects of Cre sometimes occur. However, *Cdh16^Cre^* mice is a well characterized and extensively used system with no reported off-target activity.

This work highlights an inflammatory signaling axis mediated by IL-17 and Regnase-1 and the transcription factor IκBζ (Fig 7). The gene encoding IκBζ (*Nfkbiz*) is transcriptionally induced by IL-17, thus acting as a feed-forward activator of IL-17 signaling (57, 58). IκBζ regulates transcription of *Lcn2* and *Il6* (58)*. Lcn2* is a well-known biomarker of kidney disease (27), and causes renal damage by inducing apoptosis of RTECs (26, 33, 59). This observation likely explains why we observed such a potent impact of negative regulatory mechanisms of IL-17 signaling in the development of nephritic pathology. We also observed that IL-17 synergizes with TNFa in inducing *NFKBIZ, IL6 and LCN2* in HK-2 cell lines lacking Regnase-1 (*ZC3H12A*). TNFa on its own does not activate *NFKBIZ* (60), and therefore this result reveals a mechanism by which IL-17 and TNFa together drive the expression of IκBζ and its responsive genes.

Understanding how IL-17 and its downstream signals are regulated is important from a basic science perspective and has potential translational applications. Currently, ~90 clinical trials are ongoing to test IL-17 blockade (clinicaltrials.gov: # NCT01107457; # NCT00966875; # NCT01539213; # NCT00936585; # NCT03073213). We previously showed that blocking IL-17 alleviates symptoms in mouse models of AGN (10). Antibodies against IL-23, cytokine responsible for generating pathogenic Type 17 cells, is showing promising results in clinical trials in number of inflammatory diseases (clinicaltrials.gov: #NCT01845987; # NCT01947933; # NCT04630652). Hence, future studies will need to focus on conducting pre-clinical trials to assess the efficacy of anti-IL-23, anti-IL-6 and anti-IL-1 therapies, the upstream regulators of type 17 cells development, in mouse model of AGN. Here, we have identified a novel role for the endoribonuclease Regnase-1 in restricting IL-17-mediated kidney inflammation through the novel transcription factor IκBζ. Since Regnase-1 negatively regulates pathogenic IL-17 activity in the nephritic kidney, these results may ultimately have the potential to provide new targets for therapies against AGN.

## Methods

### Mice

C57BL/6NTac (WT) mice were from Taconic Biosciences, Inc. *Il17ra*^−/−^ mice were from Amgen and bred in-house. *Il17ra^fl/fl^* mice were provided by Jay Kolls (University of Tulane). *Zc3h12a^fl/fl^* mice are under MTA from U. Central Florida. *Cdh16^Cre^* (*Ksp1.3^Cre^*) mice and B6.SJL-Ptprc (B6 CD45.1) were from The Jackson Laboratories. All the experiments used age-matched controls of both sexes, housed in specific pathogen-free conditions.

### Antibody-mediated glomerulonephritis (AGN)

Mice are immunized intraperitoneally (i.p.) with 0.2 mg of rabbit IgG (Jackson Immunoresearch) in Complete Freund’s Adjuvant (CFA) (Sigma). Controls received CFA only. Three days later, mice were injected intravenously (i.v.) with heat-inactivated rabbit anti-mouse glomerular basement membrane (GBM) serum at 5 mg/20 g body weight. Mice were sacrificed at day 7 or 14 after anti-GBM injection. Serum blood urea nitrogen (BUN) was measured using Blood Urea Nitrogen Enzymatic kit (Bioo Scientific Corp.) and creatinine with QuantiChrom Creatinine Assay kit (BioAssay Systems).

To create BM chimeras, mice were sub-lethally irradiated (9 Grey) and 24 h later 5-10 x10^6^ donor BM cells (either CD45.1 or CD45.2) were injected i.v. After 6-8 weeks, peripheral blood of recipients was tested for reconstitution with donor BM cells by flow cytometry for CD45.1 and CD45.2.

### Deletion of ZC3H12A in HK-2 cells

Knockout of *ZC3H12A* in the HK-2 cell line was performed by SYNTHEGO Company using CRISPR-Cas9. Briefly, HK-2 cells were electroporated with Cas9 and a *ZC3H12A*-specific sgRNA (Guide Sequence: GACACAUACCGUGACCUCCA). Isogenic negative control HK-2 cells were electroporated with only Cas9. Editing efficiency was evaluated 48 h post transfection. The cells were expanded to get a knockout cell pool. Clonal *ZC3H12A* knockout cell lines were generated by limiting dilution. Genomic DNA from monoclonal populations were verified by PCR and Sanger sequencing using *ZC3H12A* specific primer (F: TGACCTTGGCGTTAACCACTC; R: GTGGACCCCAAGTCTGTCAG). The sequencing results were analyzed using the Inference of CRISPR Edits (ICE) tool (https://ice.synthego.com/#/) and NCBI nucleotide blast tool.

### Immunoblot analysis

HK-2 cells (ATCC) were lysed in 1X NP-40 lysis buffer supplemented with protease inhibitor cocktail. Lysates were separated by SDS-PAGE, transferred to polyvinylidene difluoride membranes and imaged with the enhanced chemiluminescence detection system (ThermoScientific) and developed with a FluorChem E imager (ProteinSimple). Immunoblotting Abs: human/mouse anti-Regnase-1 (MAB7875, R&D Systems), human anti-β actin (ab49900, Abcam), human anti-C/EBPδ (sc-365546, Santa Cruz), human/mouse anti-C/EBPβ (sc-7962, Santa Cruz), human anti-Iκbζ (9244, Cell signaling), human/mouse anti-YY1 (sc-7341, Santa Cruz). The intensity of the protein band was measured by ImageJ software (NIH).

### Enzyme-Linked ImmunoSorbent Assay (ELISA)

HK-2 cells (ATCC) were stimulated with IL-17 and/or TNFα for different times. The supernatant was collected and ELISA was performed to detect IL-6 (8 h post stimulation, Cat # 88-7066-22, Invitrogen) and Lcn2 (24 h post stimulation, Cat # EHLCN2, Invitrogen).

### Luciferase assay

Luciferase assay was performed as previously described before (61). Briefly, IL-17 Reporter HEK 293 Cells were seeded in 12-well plates. 100 ng of Lcn2 promoter luciferase plasmid and 5 ng of Renilla luciferase plasmid were co-transfected. Cells were stimulated with 10 ng/ml of human IL-17 for 14 h and then lysed for luciferase analyze by using GloMax Navigator Microplate Luminometer.

### Histology and flow cytometry

For IHC, kidneys were fixed with 10% buffered formaldehyde and embedded in paraffin. Slices of 4 μm thick were stained with hematoxylin and eosin (H&E) or Periodic Acid Schiff (PAS) and observed on an EVOS microscope. The slides were scored blindly by an expert with more than 10 years of experience with renal histopathology, as described before (62). Briefly, severity of GN was assessed by mild to moderate increase in mesangial cellularity, thickening of the GBM, endocapillary hypercellularity and crescents formation. The tubulointerstitial inflammation is measured by assessing tubular atrophy and tubulointerstitial inflammation.

Kidneys were perfused with PBS containing EDTA (MilliporeSigma) before harvesting. Kidneys were digested at 37°C in 1 mg/ml collagenase IV (Worthington) in complete RPMI for 30 minutes, filtered through 70-mm strainers and washed twice in PBS. For flow cytometry, the following Abs are used: CD45 (30-F11), CD3 (145-2C11), CD4 (GK1.5), CD8 (53-6.7), CD45R/B220 (RA3-6B2), Ly6G (IA8), CD11b (M1/70), F4/80 (BM8) and Ly6C (HK1.4) (BD Pharmingen). Samples were acquired on BD LSRFortessa cytometer (BD Biosciences) and analyzed by FlowJo software (Tree Star).

### RNA silencing and qPCR

ON-TARGETplus SMARTpool siRNAs targeting human *NFKBIZ* were from Dharmacon. For RNA silencing, HK-2 cells were seeded overnight in antibiotic-free DMEM/F12 media (Gibco) with supplements. Transfection was performed 18 h later with 50nM siRNAs with DharmaFECT Reagent 1. Culture media was replaced after 24 h. One day later, cells were treated with human IL-17 and human TNFα (Peprotech) for 8 h.

RNA was extracted using RNeasy kits (Qiagen). Complementary DNA was synthesized by SuperScript III First Strand Kits (Thermo Fisher Scientific). Quantitative real-time PCR (qPCR) was performed with the PerfeCTa SYBR Green FastMix (Quanta BioSciences) and analyzed on an ABI 7300 real-time instrument. Primers were from QuantiTect Primer Assays (Qiagen). Expression was normalized to mouse or human *Gapdh*.

### Statistical analysis

All data are shown as Mean ± SEM. Statistical analyses were performed using unpaired T test or ANOVA through Graphpad Prism. *P < 0.05; **P < 0.01; ***P < 0.001. All experiments were performed at least twice in independent replicates.

### Study Approval

All the experiments were conducted following National Institute of Health (NIH) guidelines under protocols approved by the University of Pittsburgh IACUC (Protocol # 20087922).

## Author contributions

DL, JKK, SG, and PSB designed the experiments; DL, RB, KR, CVJ, YL, PSB performed the experiments. JKK provided conditional knock-out mouse. DL, SG, and PSB analyzed and interpreted the data and wrote the manuscript.

## Acknowledgments

This work was supported by a Rheumatology Research Foundation grant to SLG and PSB and NIH grants AI145242 (SLG, PSB), AI147383 (SLG), DK104680 (PSB), AI142354 (PSB), and R35 HL139930 (JKK). We thank PK Kollatukudy, MJ McGeachy, K Chen, S Majumder and G Trevejo-Nunez for valuable input and B. Coleman and L. Lin for assistance with mouse breeding.

## Notes

Conflict of interest statement: The authors have declared that no conflict of interest exists.

### Competing Interest Statement

The authors have declared no competing interest.

